# Variance heterogeneity genome-wide mapping for cadmium in bread wheat reveals novel genomic loci and epistatic interactions

**DOI:** 10.1101/668087

**Authors:** Waseem Hussain, Malachy Campbell, Diego Jarquin, Harkamal Walia, Gota Morota

**Author notes:** **Corresponding author**, Waseem Hussain, Department of Agronomy and Horticulture, University of Nebraska-Lincoln, Lincoln, Nebraska 68583 USA., Current address: International Rice Research Institute, Los Banos, Philippines, Gota Morota, Department of Animal and Poultry Sciences, Virginia Polytechnic Institute and State University, 175 West Campus Drive, Blacksburg, Virginia 24061 USA. **Core Ideas:** Variance-heterogeneity mapping for grain Cadmium (Cd) concentration in bread wheat was performed. Novel variance-heterogeneity loci were detected on chromosomes 2A and 2B. Loci influencing both mean and variance were identified on chromosome 5A. Identified variance-heterogeneity loci were associated with epistatic interactions. Homoeology within the vQTL on chromosomes 2A and 2B was found. ABC transporter, ATP-binding cassette transporter; Cd, cadmium; DGLM, double generalized linear model; GLM, generalized linear model; GRM, genomic relationship matrix; HGLM, Hierarchical generalized linear model; HWW, hard-red winter wheat; mQTL, mean quantitative trait loci; mvQTL, mean-variance quantitative trait loci; QTL, quantitative trait loci; ROS, reactive oxygen species; SNP, single nucleotide polymorphism; vQTL, variance heterogeneity quantitative trait loci; variance vGWAS heterogeneity genome-wide association studies.

## Abstract

Genome-wide association mapping identifies quantitative trait loci (QTL) that influence the mean differences between the marker genotypes for a given trait. While most loci influence the mean value of a trait, certain loci, known as variance heterogeneity QTL (vQTL) determine the variability of the trait instead of the mean trait value (mQTL). In the present study, we performed a variance heterogeneity genome-wide association study (vGWAS) for grain cadmium (Cd) concentration in bread wheat. We used double generalized linear model and hierarchical generalized linear model to identify vQTL associated with grain Cd. We identified novel vQTL regions on chromosomes 2A and 2B that contribute to the Cd variation and loci that affect both mean and variance heterogeneity (mvQTL) on chromosome 5A. In addition, our results demonstrated the presence of epistatic interactions between vQTL and mvQTL, which could explain variance heterogeneity. Overall, we provide novel insights into the genetic architecture of grain Cd concentration and report the first application of vGWAS in wheat. Moreover, our findings indicated that epistasis is an important mechanism underlying natural variation for grain Cd concentration.

---

Genome-wide association studies (GWAS) are routinely conducted to study the genetic basis of important traits in crops. GWAS link phenotypic variation with dense genetic marker data using a linear modeling framework (e.g., Nordborg and Weigel, 2008; Ingvarsson and Street, 2011; Huang and Han, 2014; Xiao et al., 2017). Standard GWAS approaches seek to identify marker-trait associations that influence the mean phenotypic values. However, differences in the variance between genotypes are also under genetic control (Shen et al., 2012). As a result, several recent studies have identified loci associated with differences in variance between genotypes (Cao et al., 2014; Corty et al., 2018). Such genetic variants that affect the variance heterogeneity of traits have been referred to as variance heterogeneity quantitative trait loci (vQTL) (Rönnegård and Valdar, 2011). vQTL can be detected by searching the difference in the variability between the groups of genotypes that carry alternative alleles at a particular locus (Forsberg and Carlborg, 2017). A simple example is genotypes of wheat with difference in plant height. One genotype group is homozygous for a certain allele and manifests greater variability (including both shorter and taller plants), while the second genotype group that is homozygous for the alternative allele involves plants that are similar or uniform in height. This contrast in plant height across two allelic groups leads to genetic variance heterogeneity. Note that the mean difference between the two groups does not have to be different for variance heterogeneity to arise (Fig. 1).

**Figure 1.**
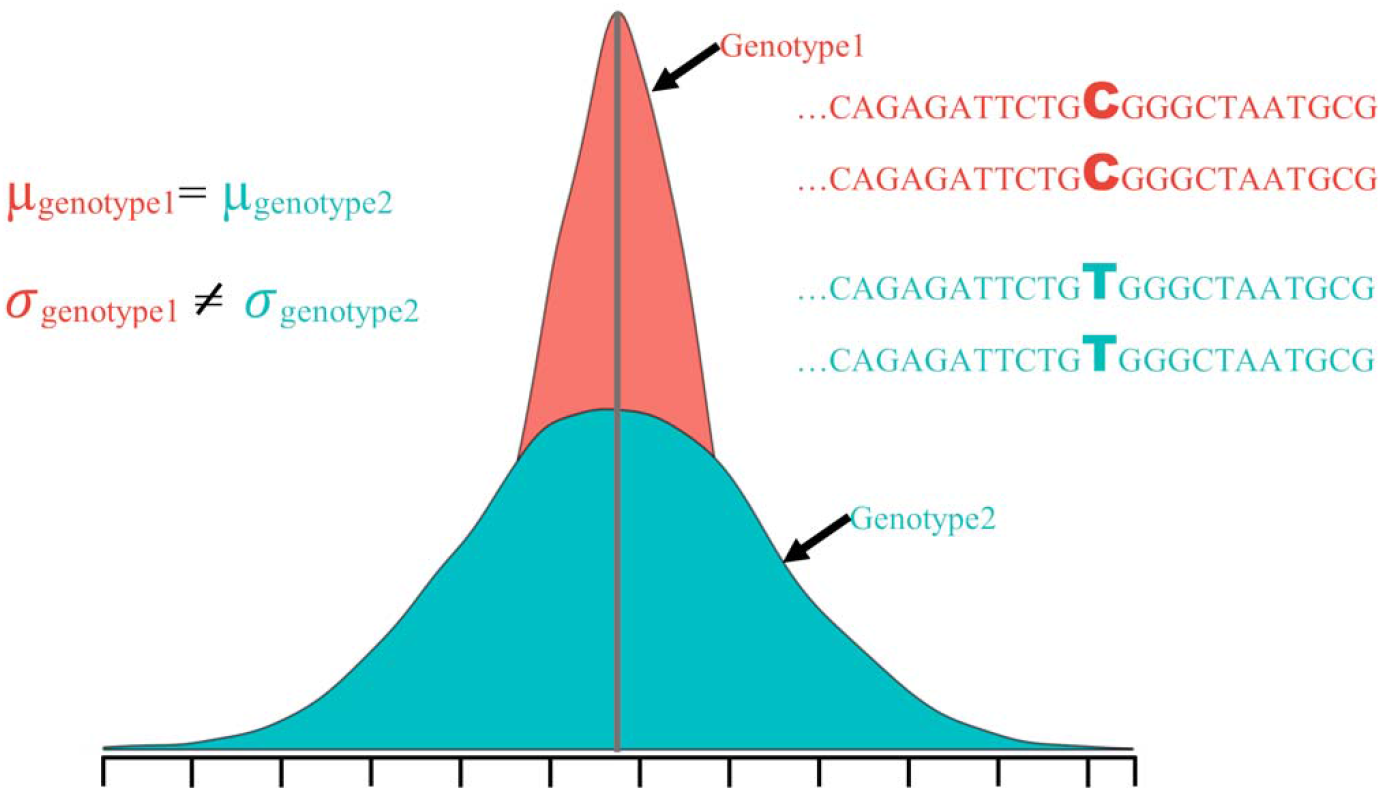
Illustration of variance heterogeneity of two genotype groups at a biallelic locus affecting the variance not the mean. Genotypes with CC allelic combination present narrow variance, whereas genotypes with TT allelic combination show greater variability. The mean difference between two genotype groups is the same as shown by the solid vertical gray line.

Variance heterogeneity-based genome-wide association studies (vGWAS) have emerged as a new approach for identifying and mapping vQTL. vQTL contribute to variability, which is undetected through standard statistical mapping (bi-parental or association) procedures (Rönnegård and Valdar, 2011; Shen et al., 2012; Forsberg and Carlborg, 2017). It has been argued that variance heterogeneity between genotypes can be partially explained by epistasis or gene-by-environment interactions (Brown et al., 2014; Forsberg and Carlborg, 2017; Young et al., 2018). Thus, vQTL can provide insights into epistasis or phenotypic plasticity (Nelson et al., 2013; Young et al., 2018). Moreover, these vGWAS frameworks can serve as tractable approaches to reduce the search space when assessing epistasis among markers (Brown et al., 2014; Wei et al., 2016). This is because we can limit the number of interacting marker pairs 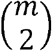 to be investigated into 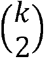, where k is the number of markers (*k* < *m*) associated with vQTL or mvQTL.

Numerous studies have reported vQTL associated with diverse phenotypes, including the tendency to left-right turning and bristles (Mackay and Lyman, 2005) and locomotor handedness (Ayroles et al., 2015) in *Drosophila*; coat color (Nachman et al., 2003), circadian activity, and exploratory behavior (Corty et al., 2018) in mice; thermotolerance (Queitsch et al., 2002), flowering time (Salomé et al., 2011), and molybdenum concentration (Shen et al., 2012; Forsberg et al., 2015) in *Arabidopsis*; litter size in swine (Sell-Kubiak et al., 2015); urinary calcium excretion in rats (Perry et al., 2012); and body mass index (Yang et al., 2012; Young et al., 2018), sero-negative rheumatoid arthritis (Wei et al., 2017), and serum urate (Topless et al., 2015) in humans. In plants, vGWAS have been limited to few species, including *Arabidopsis* (Shen et al., 2012; Forsberg et al., 2015) and maize (Kusmec et al., 2017).

Methodologically, vQTL have been detected by performing statistical tests searching for unequal variance for a quantitative trait between the marker genotypes (Rönnegård and Valdar, 2012). The most common statistical tests used to identify vQTL include Levene’s test (Paré et al., 2010), Brown-Forysthe test (Brown and Forsythe, 1974), squared residual value linear modeling (Struchalin et al., 2012), and correlation least squares test (Brown et al., 2014). However, these methods have certain drawbacks when applied to genetic data. For example, Levene’s and Brown-Forsythe tests are sensitive to deviations from normality of residuals and have an inherent inability to model continuous covariates (Rönnegård and Valdar, 2011; Dumitrascu et al., 2019).

Double generalized linear model (DGLM) has emerged as an alternative approach to model the variance heterogeneity for genetic studies (Rönnegård and Valdar, 2011). In DGLM, sample means and residuals are modelled jointly. Here, generalized linear models (GLM) are fit by including only the fixed effects in the linear predictor(s) for the mean and then the squared residuals are used to estimate the dispersion effects. It is important to correct for population structure, which can otherwise lead to spurious associations in GWAS (Patterson et al., 2006). In DGLM, population structure can be corrected by incorporating the first few principal components of a genomic relationship matrix (GRM) (Patterson et al., 2006; Price et al., 2010) as fixed covariates in the model. However, the first few principal components may not be sufficient to account for complex population structure or family relatedness (Hoffman, 2013; Sul et al., 2018). Alternatively, we can fit linear mixed models (LMM) to explicitly correct for population structure, where the whole GRM can be included to account for relationships among individuals and correct for background genotype effects. Hierarchical generalized linear model (HGLM) has been proposed as an extension of the DGLM to model random effects in the mean component (Rönnegård and Valdar, 2012; Tan et al., 2014). In HGLM, the GRM can be used to model correlated random effects and account for population structure.

We applied a vGWAS framework to examine the genetic architecture of grain cadmium (Cd) accumulation in wheat. Cd is a heavy metal that is highly toxic to human health (Menke et al., 2009). Identifying genetic variants that control low-grain Cd concentration in wheat is necessary to understand the basis for phenotypic variation in grain Cd and can help accelerate the development of low Cd wheat varieties. A recent study assessed natural variation in bread wheat grain Cd by conducting GWAS (Guttieri et al., 2015a). However, only a fraction of phenotypic variation could be explained by the top marker associations, indicating that grain Cd concentration is a complex trait that is influenced by multiple loci and/or loci with non-additive effects (Guttieri et al., 2015a). Given the genetic complexity of Cd in wheat, we hypothesized that variation in grain Cd concentration in wheat is influenced by vQTL that are likely to be involved in epistatic interactions; this would allow us to capture additional variation that is not accounted for in a standard GWAS approach.

In this study, we sought to provide additional insights into natural variation in grain Cd concentration by extending the standard GWAS to vGWAS using a hard winter wheat association mapping panel. To achieve this, we used DGLM and HGLM to perform vGWAS. Previously, Guttieri et al., (2015a) conducted the standard GWAS using this association panel and identified a single mean effect QTL (mQTL) for grain Cd concentration on chromosome 5A. In addition, we aimed to understand the basis of vQTL by searching for pairwise epistatic interactions among vQTL and mQTL. To our knowledge, the present study is the first to conduct vGWAS and identify vQTL associated with grain Cd concentration in wheat.

## MATERIALS AND METHODS

### Plant Materials and Genotyping

We analyzed a publicly available dataset comprising of phenotypes for grain mineral concentration for *n* = 299 genotyped hard-red winter wheat accessions (hereafter called as HWW association panel). The details of the study are discussed in Guttieri et al., (2015a; 2015b), and access to the data is available at http://triticeaetoolbox.org/wheat/. The data are also downloadable at https://github.com/whussain2/vGWAS/tree/master/Data. Here, we focused on grain Cd concentration (mg/kg) collected across two years (2012 and 2013) in one location (Oklahoma, USA). Briefly, the experiment was laid in an augmented incomplete block design with two replications and 15 blocks within each replication. Least square means adjusted across the replications and blocks in each year were obtained for each genotype. In this study, we averaged the least square means for each genotype across two years because of non-significant genotype x year interaction (Guttieri et al., 2015a). The association panel was genotyped using a 90K iSelect Infinium array (Wang et al., 2014b). We used a filtered marker data set consisting of *m* = 14, 731 single nucleotide polymorphism (SNP) markers from the 90K iSelect Infinium array as described by Guttieri et al., (2015a). All the SNP markers were physically anchored on the new reference genome of hexaploid wheat RefSeq v1.0 (International Wheat Genome Sequencing Consortium (IWGSC), 2018).

### Statistical Modeling

#### Genome-Wide Association Mapping

Standard GWAS or mQTL analysis based on mean differences between marker genotypes for grain Cd concentration was performed similar to Guttieri et al., (2015a) using the rrBLUP package (Endelman, 2011) in the R environment (R Core Team 2018).

#### Variance-Heterogeneity Genome-Wide Association Mapping

We used DGLM and HGLM to perform vGWAS and detect vQTL in the current study. The description of models used is given below.

#### DGLM

DGLM is a parametric approach that can be used to jointly model the mean and dispersion using a GLM framework (Smyth, 1989). The DGLM works iteratively by first fitting a linear model to estimate the mean effects (mQTL). The squared residuals are used to estimate the dispersion effects (vQTL) using GLM with a gamma-distributed response and the log link function. This process is cycled until convergence. Here, we extended the DGLM to marker-based association analysis according to Rönnegård and Valdar (2011). The mean part of DGLM was as follows:

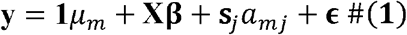

where **y** is the Cd concentration (mg/kg); **1** is the column vector of 1; μm is the intercept; **X** is *n* × 4 covariate matrix of the top four principle components (PCs) obtained by performing principal component analysis (PCA) of marker data using the SNPRelate R package (Zheng et al., 2012); **β** is the regression coefficients for the covariates; **s**_*j*_ ∈ (0,2) is the vector containing the number of reference allele at the marker *j*, *a*_*mj*_ is the effect size or allele substitution effect of the *j*th marker; and **ϵ** is the residual. We assumed

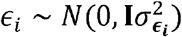

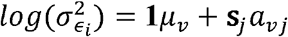

where **I** is the identity matrix; 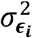 is the residual variance; and **1***μ*_*v*_ and *a*_*v*_ are the intercept and marker regression coefficients for the variance part of the model, respectively. While we fit separate effects for the mean using a standard linear model and for the variance using the squared residuals in gamma distributed GLM with a log link function, this is equivalent to modeling **y** ~ *N*(**1***μ* + **Xβ** + **s***a*_*mj*_, exp(**1***μ*_*v*_ + **s**_*j*_*a*_*vj*_) or ***ϵ*** ~ *N* (0, exp(**1***μ*_*v*_ + **s**_*j*_*a*_*vj*_)) in equation (1).

The DGLM was fitted using the dglm package (https://cran.r-project.org/web/packages/dglm/index.html) in R. SNP markers were fitted one by one, and for each marker, the effect sizes, standard errors, and p-values were obtained for the mean and dispersion components. To account for multiple testing, we determined the effective number of independent tests (Meff) using the method described by Li and Ji (2005). Subsequently, a genome-wide significance threshold level (*P* < 1.44 × 10^−5^) was determined using the following formula:

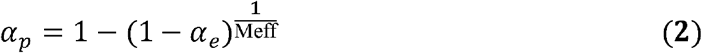

where *α*_*p*_ is the genome-wide significance threshold level, *α*_*e*_ is the desired level of significance (0.05), and Meff = 3,495.

#### HGLM

To explicitly account for population structure and kinship in GWAS, LMM have been proposed as alternative methods that allow the genetic relationships between individuals to be modeled as random effects. To perform vGWAS in the LMM framework and to identify genome-wide vQTL, we used a HGLM approach. HGLM (Lee and Nelder, 1996) is a class of GLM and is a direct extension of the DGLM that allows joint modelling of the mean and dispersion parts and introduces random effects as a linear predictor for the mean (Rönnegård and Carlborg, 2007). The mean part of HGLM was given as follows:

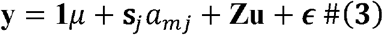

assuming that

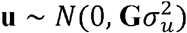

where **Z** is the incident matrix of random effects of genotypes; **u** is the vector of random effects with 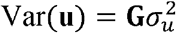; **G** is the GRM of VanRaden (2008); and 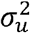 is the additive genetic variance. A log link function was used for the residual variance given by exp(***s***_*j*_ *a*_*vj*_), which is equivalent to modeling y|*a*_*mj*_, **u**, *a*_*vj*_ ~ *N*(**s**_*j*_*a*_*mj*_ + **Zu**, exp(**s**_*j*_*a*_*vj*_)).

We fitted HGLM using the hglm R package (Rönnegård et al., 2010b). We reformulated the term **Zu** as **Z**^*^ **u**^*^, where 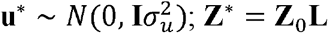; **L** is the Cholesky factorization of the **G** matrix; and **Z**_0_ is the identity matrix (Rönnegård et al., 2010a). Markers treated as fixed effects were fit one by one, and for each marker, the effect sizes, standard errors, and p-values were obtained for the mean and dispersion components. The genome-wide significance threshold level was derived as described in the DGLM analysis. Circular Manhattan and quantile-quantile (QQ) plots were created using the CMplot R package (https://github.com/YinLiLin/R-CMplot).

#### Epistasis Analysis

We investigated the extent of epistasis that was manifested through variance heterogeneity. All the possible pairwise interaction analyses for markers that were associated with grain Cd concentration were performed using the following two markers at a time epistatic model:

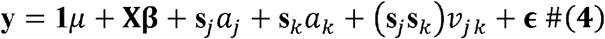

where **y** is the vector of Cd concentration (mg/kg); **x** is the incident matrix for the first four PCs; **β** is the regression coefficients for the PCs; **s**_*j*_ and **s**_*k*_ are SNP codes for the *j*th and *k*th markers, respectively; *a*_*j*_ and *a*_*k*_ are the additive effects of the markers *j* and *k*, respectively; and *v*_*jk*_ is the additive × additive epistatic effect of the *j*th and *k*th markers. We used Bonferroni correction to account for the multiple testing. The threshold of −log10(0.05/325) = −log10 (1.54 × 10^−04^) = 3.8 was used to declare the significance of interaction effects.

#### Homoeology and Candidate Gene Analysis

Homoeologous gene construction was performed as per procedure described by (Santantonio et al., 2019). Briefly, the annotated coding sequences within the 2A vQTL were aligned back onto themselves using the IWGSC RefSeq v.1.0 coupled with BLAST tool in Ensemble Plants browser (Bolser et al., 2017). For candidate gene identification for the SNP markers associated with variance heterogeneity, we used Ensembl Plants browser to retrieve the candidate genes and functional annotations (http://plants.ensembl.org/Triticum_aestivum/Info/Index) and the wheat RefSeq v1.0 annotations (International Wheat Genome Sequencing Consortium (IWGSC) et al., 2018) available at https://wheat-urgi.versailles.inra.fr/Seq-Repository/Annotations. For candidate gene analysis, we first determined the positions of significant SNP markers, and the interval was defined as the distance between the lowest and highest markers based on the position of SNP. For example, if the position of the lowest SNP and highest SNP was 715,333,165 bp and 717,146,211 bp in the vQTL region on chromosome 2A, we defined 2A as the 715,333,165-717,146,211 interval for candidate gene identification. After defining the interval for the 2A (2A: 715,333,165-717,146,211) and 2B (2B: 691,780,716-701,097,263 bp) regions, we explored the intervals using Ensembl Plants browser and extracted the Gene IDs within these intervals. The Gene IDs within the defined interval on chromosomes 2A and 2B were analyzed using the IWGSC RefSeq v.1.0 (International Wheat Genome Sequencing Consortium (IWGSC) et al., 2018) integrated genome annotations to obtain the predicted genes and functional annotations.

## RESULTS

### Variance Heterogeneity GWAS Provide Additional Insights into Natural Variation in Grain Cd

Although grain Cd concentration is a highly heritable trait, recent GWAS revealed that significant loci can only explain a fraction of the variation for this trait (Guttieri et al., 2015a). We found the single genomic region on chromosome 5A affecting the grain Cd concentration (Fig. 2) from the standard GWAS analysis confirming the results of Guttieri et al., (2015). The DGLM and HGLM approaches were used to detect vQTL while controlling for population structure. The population structure based on PCA of the HWW association panel is given in Supplemental File S1: Figure S1. QQ plots (Supplemental File S1: Figure S2) show that both DGLM and HGLM had adequate control of population structure and effective control of false positives.

**Figure 2:**
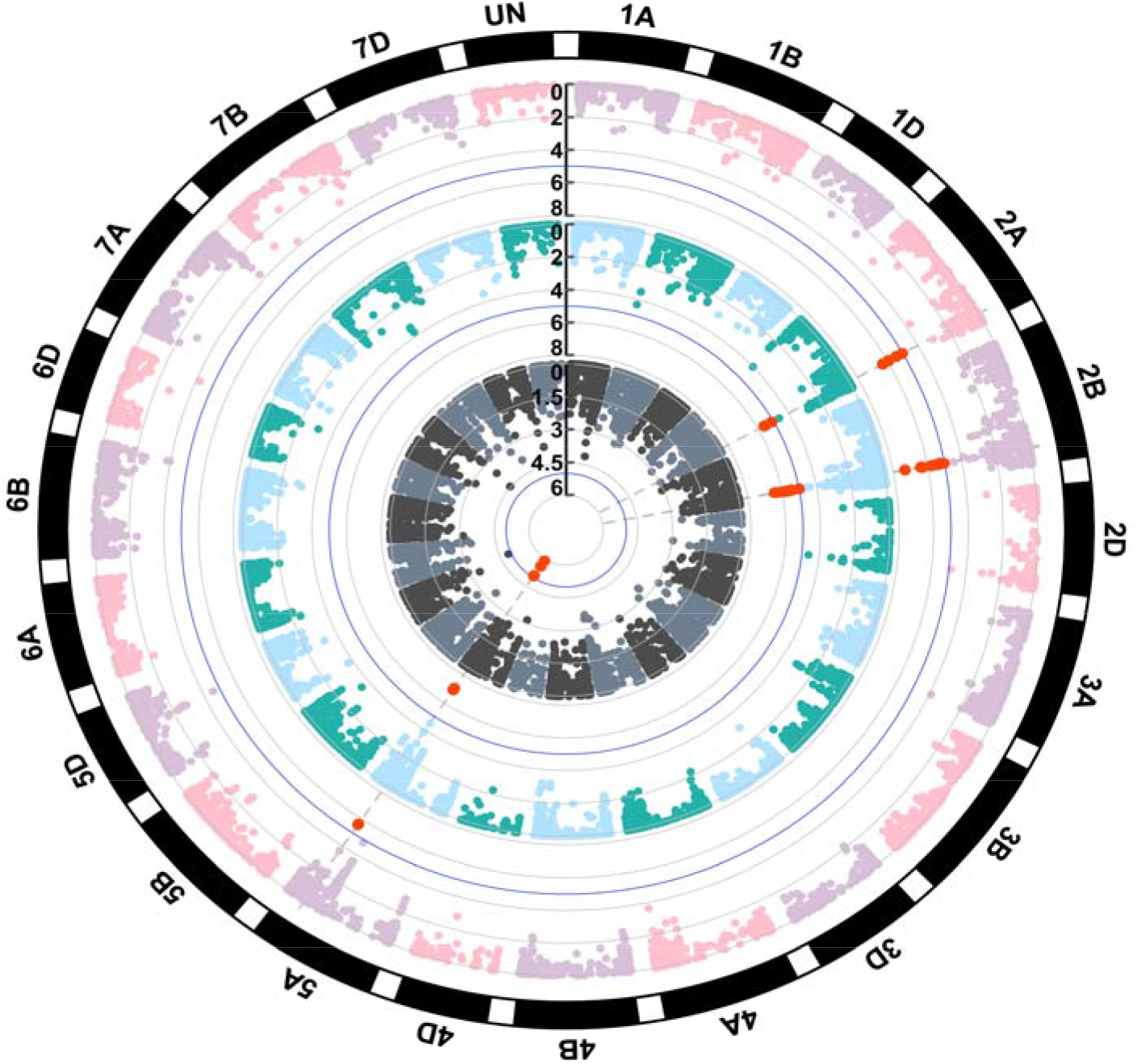
Circular Manhattan plot of standard genome-wide association studies (GWAS) based on mean differences (inner), and variance GWAS using double generalized linear model (middle) and hierarchical generalized linear model (outer) for grain cadmium concentration in the hard-red winter wheat association panel. The red dots represent the significant markers associated with either mean or variance heterogeneity quantitative trait loci. The blue line in each circular plot shows the cutoff for the statistical significance. The P-values in −log_10_ scale are given in black vertical line.

We classified the QTL into the following categories: mQTL, which contributes to difference in the means between marker genotypes; vQTL, which influences the variability between the genotypes; and mean-variance QTL (mvQTL), which contributes to differences in both the mean and variance between the genotypes.

Based on the DGLM, we identified two vQTL associated with the variance heterogeneity of Cd concentration. One vQTL on 2A contained four SNP markers, and one vQTL on 2B contained 17 SNP markers (Fig. 2 and Supplemental File S1: Table S1). The four SNP markers associated with the vQTL region on the chromosome 2A region spanned the physical distance of 1.81 Mb; all SNP markers were located within the 1,000 bp linkage disequilibrium (LD) block (Supplemental File S1: Figure S3). The vQTL region on 2B associated with 17 SNP markers spanned the physical distance of 9.32 Mb, and the SNP markers were located within four LD blocks of sizes 0, 1, 1, and 204 kb (Supplemental File S1: Figure S4).

In addition, we identified a single mvQTL (containing four SNP markers) associated with both mean and variance heterogeneity on chromosome 5A (Fig. 2 and Table S2). The markers associated with mvQTL on chromosome 5A were identical to those obtained in the original GWAS analysis according to Guttieri et al., (2015), indicating that this region affects both the mean and the variance heterogeneity (Supplemental File S1: Figure S5). Moreover, these results showed that DGLM serves as an accurate framework to jointly detect mean and variance QTL and provides additional insights into phenotypic variation that would otherwise not be captured by standard GWAS.

The HGLM analysis revealed the same results as those obtained using DGLM and showed identical vQTL on chromosomes 2A and 2B and mvQTL on chromosome 5A associated with variance heterogeneity of Cd concentration (Fig. 2 and Supplemental File S1: Table S1). Further, we observed a potential vQTL region on 2D from the DGLM and HGLM analyses. This region was slightly below the significance threshold level but may have an implication on Cd variation given that the allopolyploid nature of wheat and the role of homoeologous gene sets on phenotypic variation (Borrill et al., 2019).

### Variance Heterogeneity Loci can be Partially Explained by Epistasis

We investigated all significant markers (25 markers) associated with mvQTL on chromosome 5A and vQTL on chromosomes 2A and 2B and explored all possible pairwise additive × additive epistatic interactions. We detected significant additive × additive interactions between the markers (Fig. 3). The interaction was more evident between mvQTL on chromosome 5A and vQTL on chromosomes 2A and 2B. Specifically, all the markers associated with the 5A mvQTL region revealed highly significant interactions with all the markers associated with the 2A and 2B vQTL regions. Interactions between vQTL on chromosomes 2A and 2B were also observed; however, the interactions were less evident, and only a few markers within these regions showed statistically significant interactions. Taken together, these results suggested that the vQTL and mvQTL may be manifested because of pairwise epistatic interactions.

**Figure 3:**
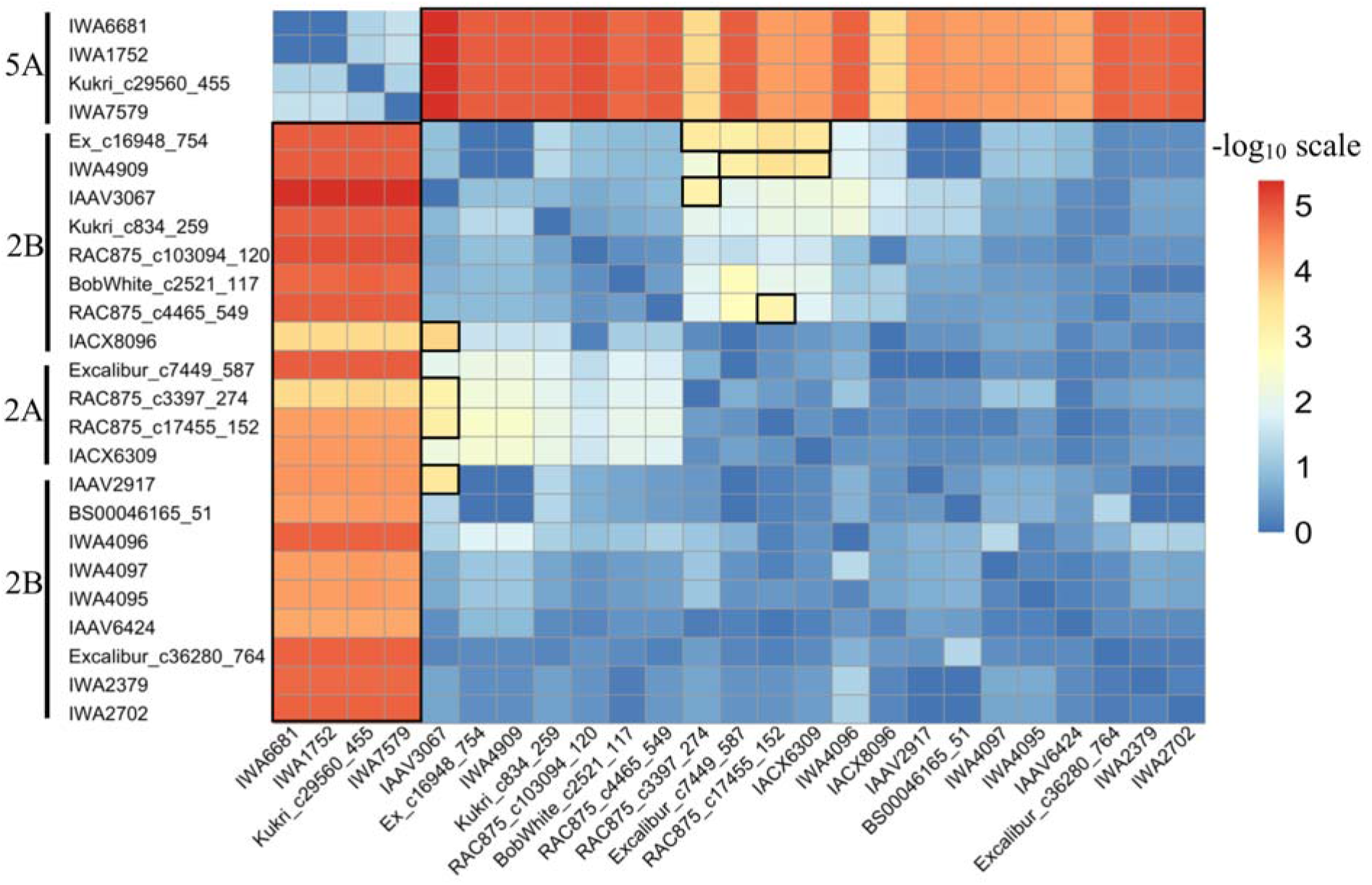
Heat map showing all possible pairwise epistatic interactions between the markers associated with vQTL on chromosomes 2A and 2B or mvQTL on chromosome 5A. Chromosome information of each marker is given on the left side. The heat map is sorted and color coded based on −log_10_ (p-value) scale with the legend given on right side. Interactions that are significant (−log_10_ > 3.8) are color coded as red or orange in color and outlined in black box.

### Homoeology and Candidate Genes

Homoeology analysis between the defined regions on chromosomes 2A and 2B resulted in 22 homoeologous gene sets, consisting of 21 triplicates and only one duplicate gene set. Additional details on the homoeologous gene sets can be found in Supplemental File S2. As compared to the total number of candidate genes equal to 39 within the 1.18 Mb 2A region, 22 (58%) were homoeologous across the three genomes. Based on the annotations for the 22 homoeologous gene sets, a few of the genes encoded homeobox-leucine zipper family protein, plant peroxidase, and glycosyltransferase, which have been associated with the genetic regulation of minerals in plants (Whitt et al., 2018). For example, homeodomain-leucine zipper family protein has been functionally associated with Cd tolerance by regulating the expression of metal transporters *OsHMA2* and *OsHMA3* in rice (Ding et al., 2018; Yu et al., 2019). These genes have been found to play important roles in loading Cd onto the xylem and root-to-shoot translocation of Cd in rice. In plants, response to heavy metals involves the accumulation of reactive oxygen species (ROS) that damage DNA and cellular machinery (Kumari et al., 2008; Rascio and Navari-Izzo, 2011). In *Arabidopsis*, the peroxidase genes *At2g35380*, *PER20*, and *At2g18150* have been found to be associated with Cd responses by affecting the lignin biosynthesis in root cells under high Cd stress (Chen and Kao, 1995; van de Mortel et al., 2008). Full list of candidate genes within the 2A and 2B region, and within the homoeologous gene sets is in Supplemental File S2. These results clearly indicate that most of the genes with vQTL regions are redundant across the genomes and may have significant role in the genetic regulation of grain Cd concentration in wheat. However, we contend that further investigation of these regions using dense markers and increased sample size is necessary to fine-map the QTL and validate potential candidate genes underlying these loci and also the role of gene redundancy in generating phenotypic variation.

## DISCUSSION

In the present study, we explored the genetic variants affecting variance heterogeneity of Cd. Given the complexity of genetic regulation of Cd in wheat (Guttieri et al., 2015a) and the influence of epistatic interactions, we anticipated that partial genetic regulation of Cd in wheat can be detected using methods that have been developed to identify vQTL. As reported by Rönnegård and Valdar, (2012), a potential explanation for variance-controlling QTL is epistatic interactions that are unspecified in the model. Herein, we utilized two approaches, namely, DGLM and HGLM, to detect vQTL and mvQTL associated with grain Cd concentration in wheat.

The DGLM framework is a powerful approach for vGWAS analysis. However, in DGLM, GLM is fit by including only the fixed effects in the linear predictor of mean and dispersion. Therefore, by using the DGLM approach, population structure can only be accounted for by using the first few PCs obtained from the SNP matrix; however, this may not completely account for complex population structure and family relationships (Price et al., 2010). We hypothesized that the use of random effects to model the mean component can better account for population structure and reduce spurious associations. In this approach, a random additive genetic effect is introduced to the mean component of the model that accounts for population structure and cryptic relatedness between accessions. Therefore, we performed vGWAS analysis using HGLM. Interestingly, both DGLM and HGLM approaches were effective in identifying the genetic variants controlling variability of Cd, suggesting that the loci detected with the DGLM approach are likely to be true QTL rather than artifacts from population structure. The impact of population structure on the power of DGLM and HGLM remains to be explored; further examination is warranted.

In the literature, it has been argued that variance heterogeneity can also arise by a simple mean variance relationship, which does not have biological significance (Young et al., 2018). To rule out the role of the mean-variance function in generating variance heterogeneity, we plotted the estimated effects of the top three significant associated vQTL markers at the alternate genotypes and observed that the means of all the markers were the same (Fig. 4), indicating that the effect of SNP on variance heterogeneity was not due to the consequences of mean-variance function but likely due to the genetic effects (Yang et al., 2012).

**Figure 4:**
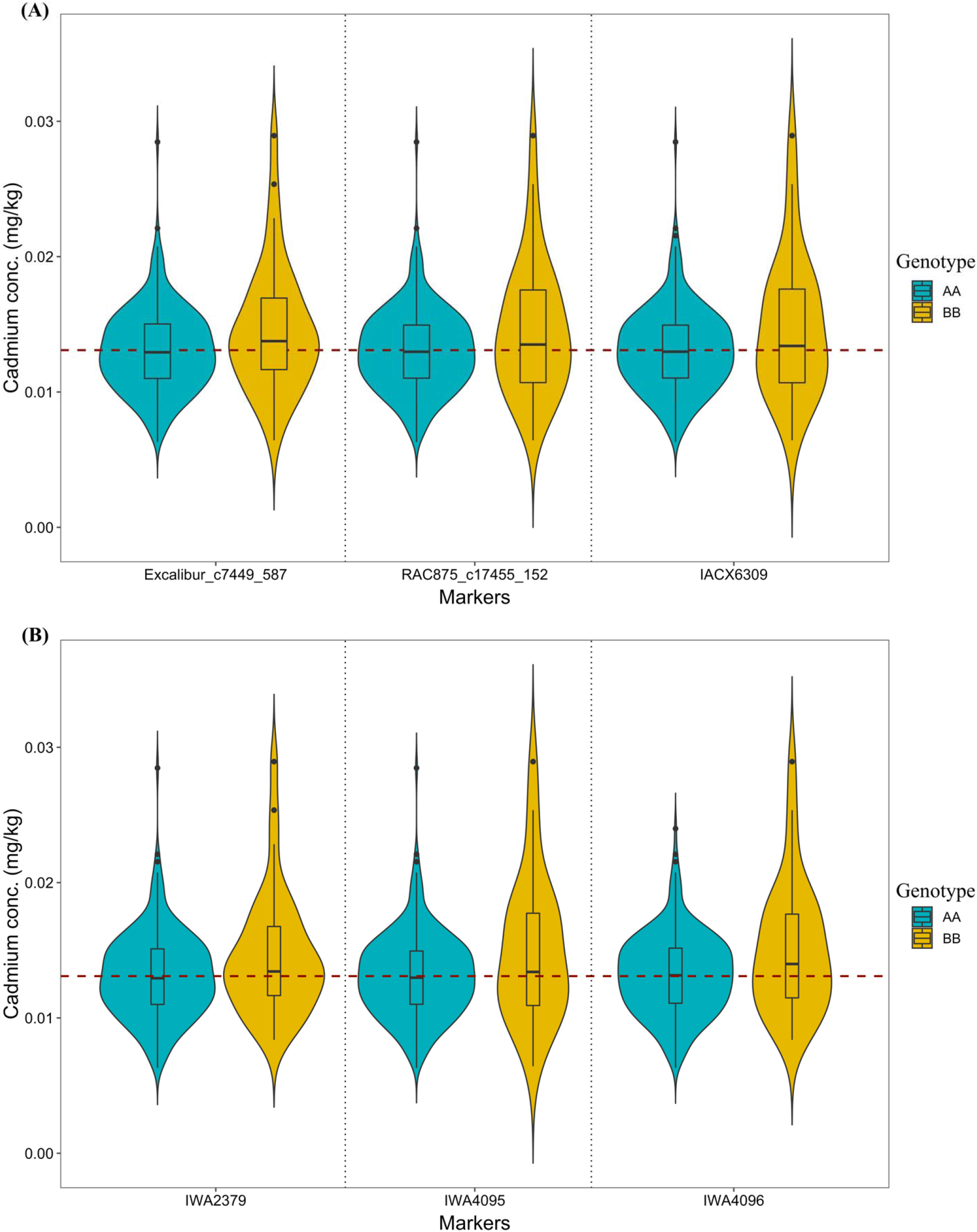
Violin plot showing the differences in the mean and variance of grain cadmium concentration with alternative marker genotype groups coded as AA and BB for the top three significant markers associated with vQTL on (A) chromosome 2A and (B) chromosome 2B. The mean of marker genotypes AA and BB are connected by red dotted line.

Further, variance heterogeneity can also be observed in a population when two or more alleles having different effects on the phenotype are in high LD (Cao et al., 2014; Wang et al., 2014a; Forsberg and Carlborg, 2017). To rule out the possibility of LD as a source for variance heterogeneity in grain Cd in this population, we suggest the use of high-density markers and larger sample size to identify the actual functional alleles associated with Cd, their LD patterns, and their effects on the Cd phenotype (Struchalin et al., 2010; Forsberg and Carlborg, 2017).

In QTL studies, variance heterogeneity arises because of various underlying mechanisms, such as epistatic interactions (Struchalin et al., 2010; Shen et al., 2012; Nelson et al., 2013). Epistasis gives rise to variance heterogeneity when the different allele combinations at one locus change the effect of the other loci in the genome, as shown in one pair of interacting markers (Fig. 5). Hence, identifying the loci affecting variance heterogeneity through vGWAS means that the loci are likely to be involved in epistatic interactions. To validate this assumption and investigate whether epistasis can explain the identified vQTL and mvQTL in this study, we analyzed all possible pairwise interactions between the associated markers. We detected significant epistatic interactions between the associated markers (Fig. 2), which can explain the existence of variance heterogeneity in the genotypes. Additionally, identifying vQTL through vGWAS serves as an effective way to restrict the search space when detecting epistatic QTL. Thus, with the vGWAS approach, many of the requirements necessary for conventional epistasis mapping can be avoided (e.g., large sample size and extensive multiple testing corrections that reduce power). However, Forsberg and Carlborg (2017) empirically showed that the presence of variance heterogeneity does not always guarantee the presence of epistatic interactions that contribute to the total variation of the trait; therefore, the results should be interpreted carefully when multi-locus interactions are involved.

**Figure 5:**
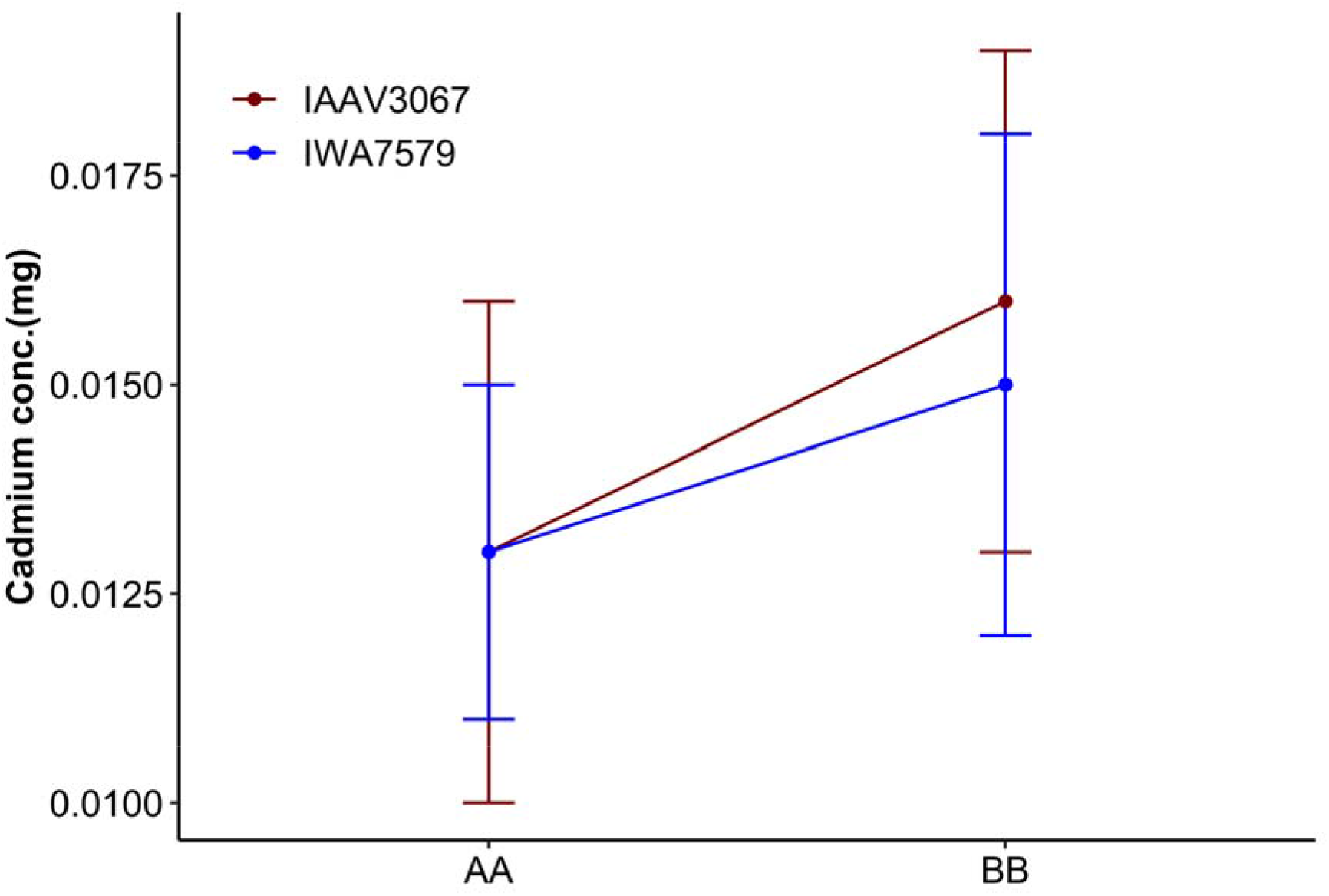
Epistatic interaction plot between marker pair IAAV3067 (shown in dark-red color) and IWA7579 (shown in blue color) on chromosomes 5A (mvQTL) and 2B (vQTL). The y-axi shows the phenotypic value of cadmium concentration (mg). AA and BB represent the alternat genotypes at the particular marker. Plotted points indicate two-locus genotype means ± standard deviations for the two loci represented by error bars. Large difference in the mean value of cadmium concentrations at BB genotype compared to no difference in the mean value of cadmium concentrations at AA genotype indicates the presence of interaction between the two markers.

The genomic regions on chromosomes 2A and 2B associated with variance heterogeneity revealed homoeologous gene sets with 58% genes revealing the gene redundancy mostly present as three functional homoeologous copies (triplicated). This also indicates that genetic complexity of Cd phenotype is not only controlled by multiple genes but may be affected by the multiple homoeologs of the individual genes which warrants further investigation. Presence of multiple copies of homoeologous genes may have consequence on phenotypic variation due to dosage effects and or functional redundancy (Borrill et al., 2019). Dosage effect, in which the phenotypic variation is amplified by the addition of each gene copies can act additively (e.g., genes controlling grain protein content (Avni et al., 2014) and grain size (Wang et al., 2018)) or non-additively (e.g., genes controlling amylopectin content in wheat (Kim et al., 2003)). Non-additive variation between homoeologous gene has been shown to be an important source of variation in wheat. However, its relative contribution across the wheat genome as compared to non-syntenic regions was proportionately less (Santantonio et al., 2019). This is in agreement with our results because we observed interactions among the homoeologous genomic regions on chromosomes 2A and 2B. However, this homoeologous gene interactions was less evident as compared to two-way interactions found between non-syntenic vQTL regions on 2A and 2B with the mvQTL region on 5A. The nature and functional role of homoeologous gene sets within the vQTL region on 2A and 2B is not clear. However, it is increasingly feasible in wheat to examine the effects of gene redundancy and explore the contribution of homoeologous genes in generating phenotypic variation (Wang et al., 2018).

## Conclusion

We showed the potential of vGWAS for dissecting the genetic architecture of complex traits and identifying novel genomic regions influencing variance heterogeneity in wheat. We provided evidence that the vQTL contribute to natural variation in grain Cd concentration through non-additive genetic effects. This is particularly evidenced by epistatic interactions between mvQTL on chromosome 5A and vQTL on chromosomes 2A and 2B.

## Acknowledgements

This work was supported by the National Science Foundation under Grant Number 1736192 to H.W. and G.M. Data analysis was performed using the Holland Computing Center computational resources at the University of Nebraska-Lincoln.

## Supplemental Materials

Supplemental File S1 contains Table S1 and Figures S1-S5.

Table S1: Single nucleotide polymorphism markers associated with variance heterogeneity of cadmium concentration in the hard-red winter wheat association panel.

Table S2: Single nucleotide polymorphism markers associated with the mean of cadmium concentration in the hard-red winter wheat association panel.

Figure S1: Principal component analysis of the population structure in the hard-red winter wheat association panel. The different colors represent the sub-populations of red wheat and winter wheat.

Figure S2: Quantile-quantile (QQ) plot of the outputs for the double generalized linear model and the hierarchical generalized linear model shown in the Manhattan plot.

Figure S3: Linkage disequilibrium block and annotated genes on chromosome 2A.

Figure S4: Linkage disequilibrium blocks and annotated genes on chromosome 2B.

Figure S5: Violin plot showing the differences in the mean and variance of Cadmium concentration with alternative marker allele groups.

Supplemental File S2: A list of candidate genes and homoeologous gene sets associated with the vQTL on chromosomes 2A and 2B.

## Data Availability

The wheat phenotypic and genotypic data can be downloaded from (http://triticeaetoolbox.org/wheat/) and also available on the GitHub repository https://github.com/whussain2/vGWAS. The R code used for the analysis is available on the GitHub repository https://github.com/whussain2/vGWAS.

## Conflict of interest

The authors declare there are no competing interests.

## Author’s Contributions

W.H. and G.M. conceived the study. W.H. performed the data analysis and drafted the manuscript. D.J. helped the data analysis. M.C., D.J., H.W., and G.M. revised the manuscript. G.M. supervised and directed the study. All authors read and approved the manuscript.

## Notes

#### Summary of Updates

The version has been revised to incorporate additional analysis on homoelogou nature of vQTL regions as suggested by the reviewers. Candidate gene analysis has been de-emphasized from the main content. Content of the manuscript has been throughly edited and updated, figure 1 is newly added figure and all other figures have been modified.

https://github.com/whussain2/vGWAS

