## Supplementary File S1 for "Variance heterogeneity genome-wide mapping for cadmium in bread wheat reveals novel genomic loci and epistatic interactions"

**Supplemental Material**

**Tables**

| **Table S1**: Single nucleotide polymorphism markers associated with variance heterogeneity of cadmium concentration in the hard-red winter wheat association panel. | | | | | | | | | | |
| --- | --- | --- | --- | --- | --- | --- | --- | --- | --- | --- |
| SNP: | Chr^1^ | P-values | | Alleles | Pos(bp)^2^ | MJF^3^ | MAF^4^ | N^5^ | AA^6^ | BB^6^ |
|  |  | DGLM | HGLM |  |  |  |  |  |  |  |
| IWA4096 | 2B | 6.50E-07 | 4.80E-08 | G/C | 696677568 | 0.89 | 0.10 | 299 | 268 | 31 |
| IWA4097 | 2B | 3.46E-06 | 4.79E-07 | T/C | 696678792 | 0.91 | 0.08 | 299 | 274 | 25 |
| IWA4095 | 2B | 3.71E-06 | 4.89E-07 | A/G | 696677327 | 0.90 | 0.09 | 299 | 270 | 29 |
| Excalibur_c36280_764 | 2B | 2.49E-06 | 4.83E-07 | T/G | 697512104 | 0.89 | 0.10 | 299 | 267 | 32 |
| IWA2379 | 2B | 2.69E-06 | 4.83E-07 | T/C | 701097263 | 0.87 | 0.12 | 299 | 261 | 38 |
| BS00046164_51 | 2B | 3.13E-06 | 4.22E-07 | A/C | 697510373 | 0.90 | 0.09 | 299 | 270 | 29 |
| IWA2702 | 2B | 2.92E-06 | 4.59E-07 | T/C | 700456554 | 0.89 | 0.10 | 299 | 267 | 32 |
| IAAV6424 | 2B | 3.67E-07 | 5.00E-07 | T/C | 691780716 | 0.87 | 0.12 | 299 | 261 | 38 |
| IAAV3067 | 2B | 1.36E-06 | 5.56E-06 | C/G | 699173434 | 0.90 | 0.09 | 299 | 271 | 28 |
| Kukri_c834_259 | 2B | 1.20E-06 | 4.89E-06 | T/C | 698968658 | 0.91 | 0.08 | 299 | 273 | 26 |
| IAAV2917 | 2B | 9.87E-06 | 5.64E-06 | A/G | 697511682 | 0.88 | 0.11 | 299 | 266 | 33 |
| BobWhite_c2521_117 | 2B | 1.31E-06 | 3.92E-06 | A/G | 699106309 | 0.87 | 0.12 | 299 | 262 | 37 |
| Ex_c16948_754 | 2B | 1.80E-06 | 4.15E-06 | A/G | 699827018 | 0.91 | 0.08 | 299 | 274 | 25 |
| IWA4909 | 2B | 1.80E-06 | 4.15E-06 | A/G | 700207481 | 0.91 | 0.08 | 299 | 274 | 25 |
| RAC875_c4465_549 | 2B | 2.26E-06 | 5.28E-06 | A/G | 699108736 | 0.90 | 0.09 | 299 | 270 | 29 |
| IACX8096 | 2B | 3.22E-06 | 5.57E-06 | G/C | 691780864 | 0.86 | 0.13 | 299 | 260 | 39 |
| RAC875_c103094_120 | 2B | 2.43E-06 | 9.05E-06 | A/C | 698303834 | 0.90 | 0.09 | 299 | 271 | 28 |
| Excalibur_c7449_587 | 2A | 2.50E-06 | 4.18E-06 | T/G | 717146211 | 0.87 | 0.12 | 299 | 262 | 37 |
| RAC875_c17455_152 | 2A | 2.93E-06 | 4.64E-07 | T/C | 715333875 | 0.90 | 0.09 | 299 | 271 | 28 |
| IACX6309 | 2A | 7.77E-06 | 4.85E-06 | A/G | 715333830 | 0.89 | 0.10 | 299 | 267 | 32 |
| RAC875_c3397_274 | 2A | 2.26E-06 | 1.20E-06 | T/G | 715333165 | 0.89 | 0.10 | 299 | 267 | 32 |

SNP: Single nucleotide polymorphism; Chr: Chromosome; Pos(bp): Physical position of marker on the wheat reference genome; MJF: Major allele frequency; MAF: Minor allele frequency; N: Total number of genotypes; AA and BB: Alternate genotypes at a specific SNP marker

| **Table S2**: Single nucleotide polymorphism markers associated with the mean of cadmium concentration in the hard-red winter wheat association panel. | | | | | | | | | |
| --- | --- | --- | --- | --- | --- | --- | --- | --- | --- |
| SNP | Chr | P-values | Alleles | Pos(bp) | MJF | MAF | N | AA | BB |
| Kukri_c29560_455 | 5A | 8.82E-08 | T/C | 581302317 | 0.70 | 0.29 | 299 | 211 | 88 |
| IWA7579 | 5A | 9.47E-08 | A/G | 581108961 | 0.70 | 0.29 | 299 | 212 | 87 |
| IWA6681 | 5A | 1.13E-07 | T/G | 580939178 | 0.70 | 0.29 | 299 | 212 | 87 |
| IWA1752 | 5A | 1.13E-07 | A/G | 580946575 | 0.70 | 0.29 | 299 | 210 | 89 |

SNP: Single nucleotide polymorphism; Chr: Chromosome; Pos(bp): Physical position of marker on the wheat reference genome; MJF: Major allele frequency; MAF: Minor allele frequency; N: Total number of genotypes; AA and BB: Alternate genotypes at a specific SNP marker

**Figures**

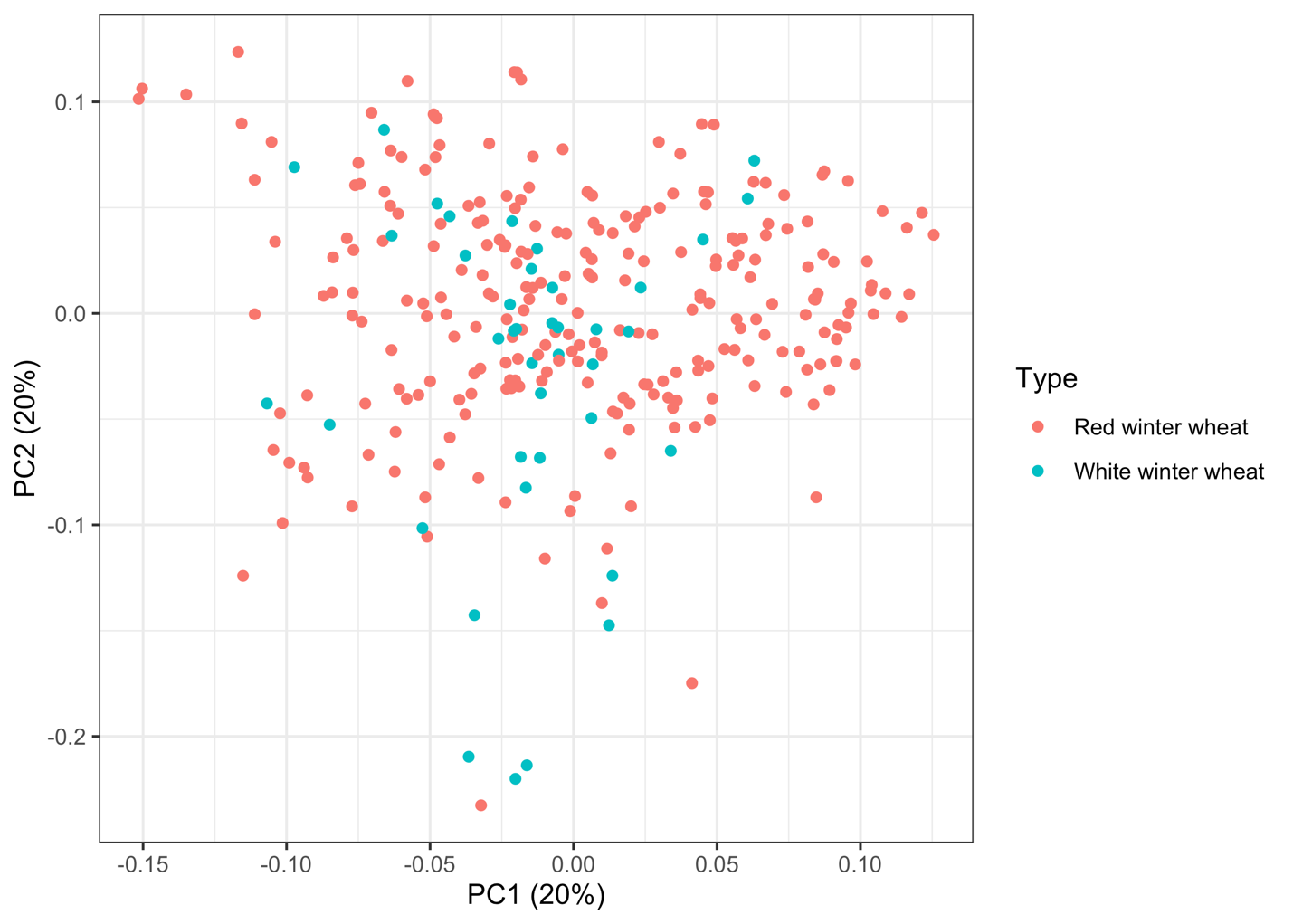

Figure S1: Principal component analysis of the population structure in the hard-red winter wheat association panel. The different colors represent the sub-populations of red wheat and winter wheat.

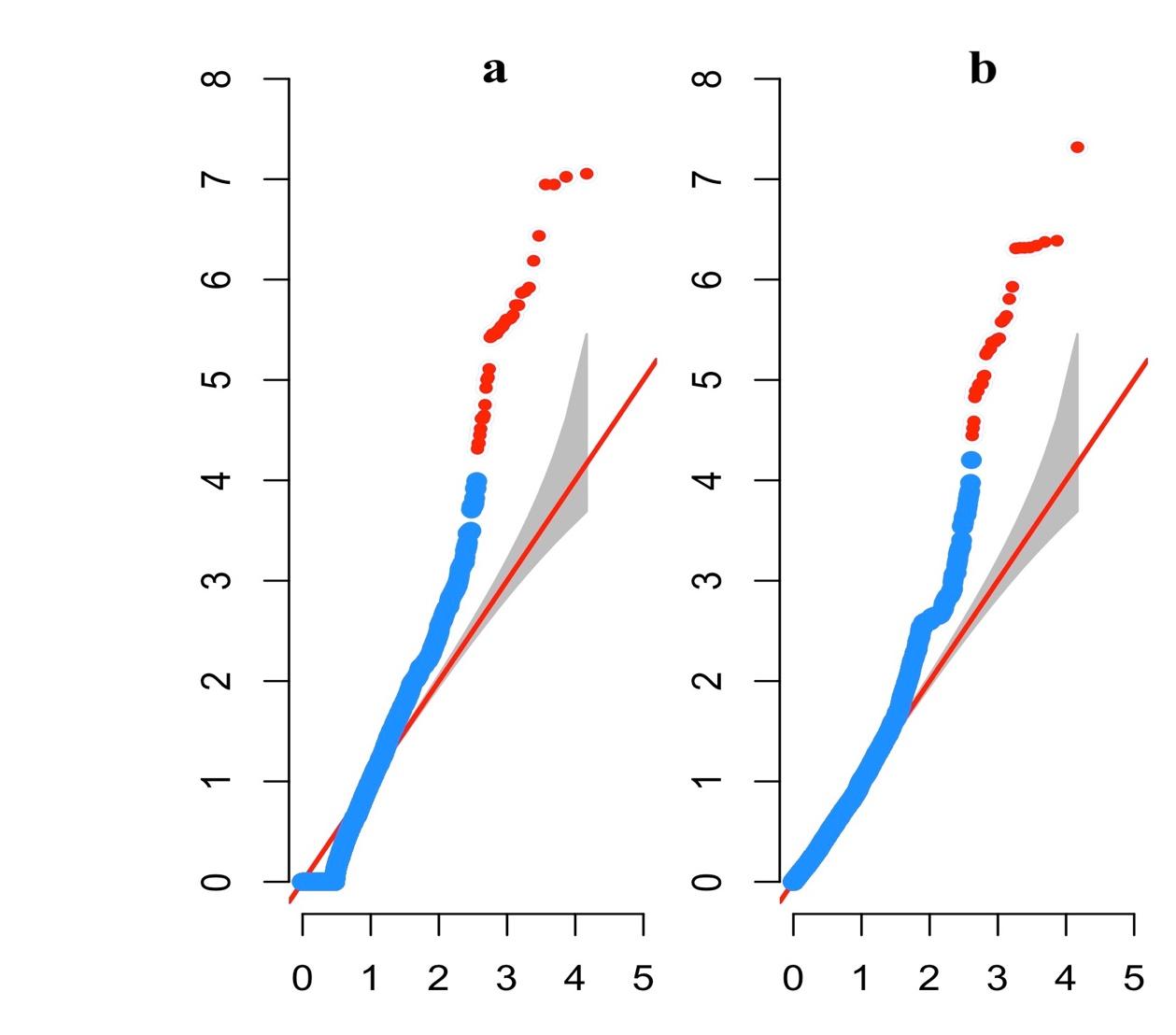

**Figure S2**: Quantile-quantile (QQ) plot from (a) double generalized linear model and (b) hierarchical generalized linear model shown in the circular Manhattan plot. The QQ plot shows the deviation of observed p-values (blue color) from expected p-values (red line) and are sorted from largest to smallest in the plot. Some markers have -log_10_ (p-value) equal to zero because of the same residual variance across the allelic groups for a marker.

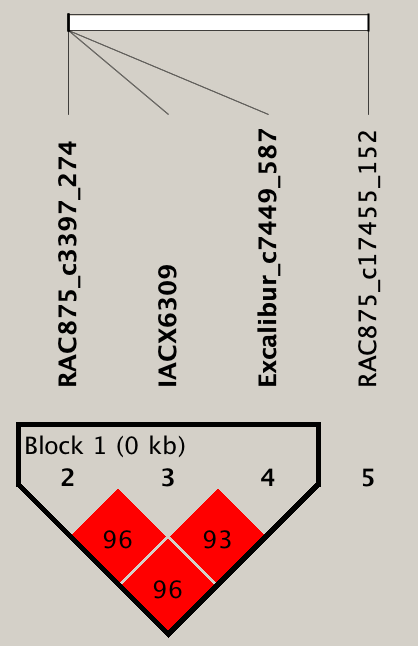

**Figure S3**: Linkage disequilibrium block on chromosome 2A in the vQTL region of 715,333,165 to 717,146,211 bp. A single block of less than 1,000 bp within the vQTL region.

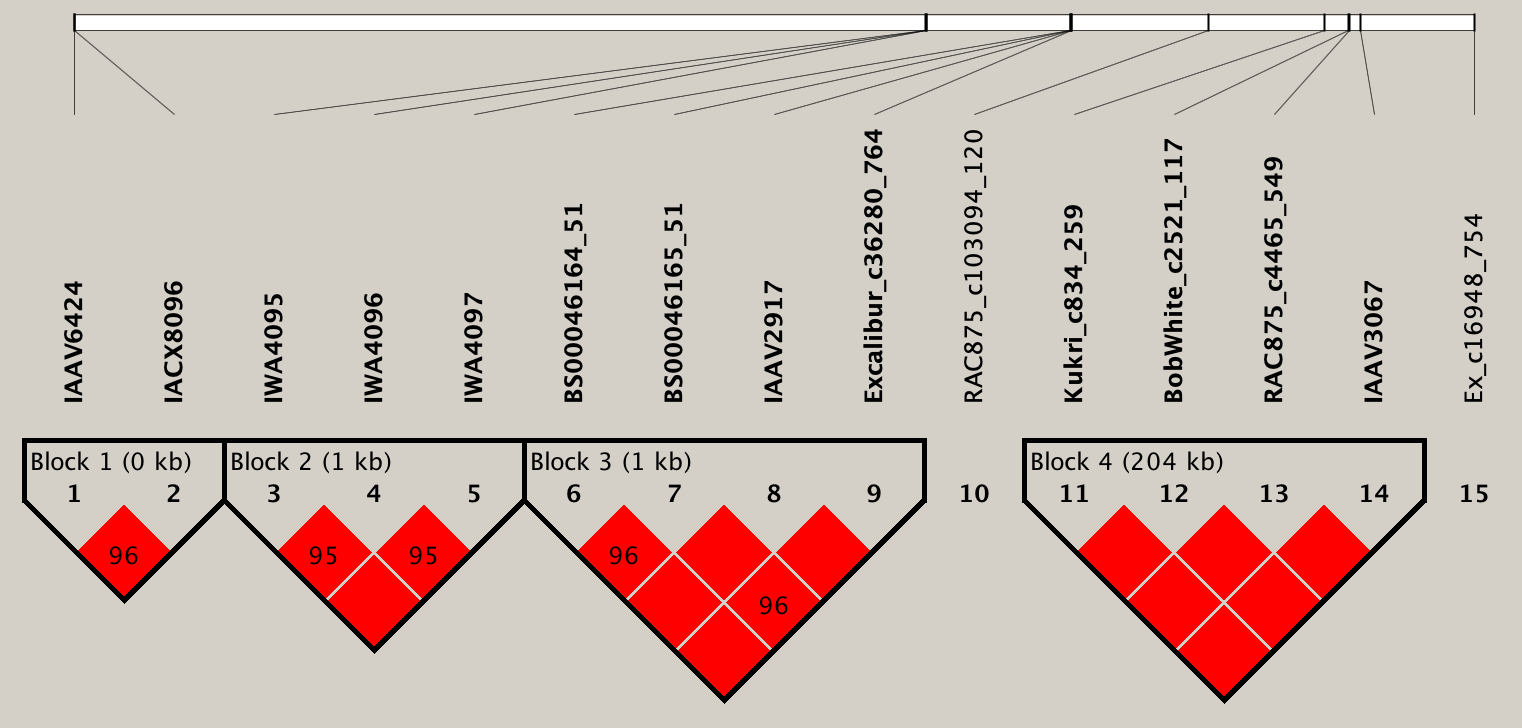

**Figure S4**: Linkage disequilibrium blocks on chromosome 2B in the vQTL region of 691,780,716 to 699,827,018 bp. A four linkage disequilibrium blocks of 0 (less than 1,000 bp), 1, 1, and 204 kb within the vQTL region.

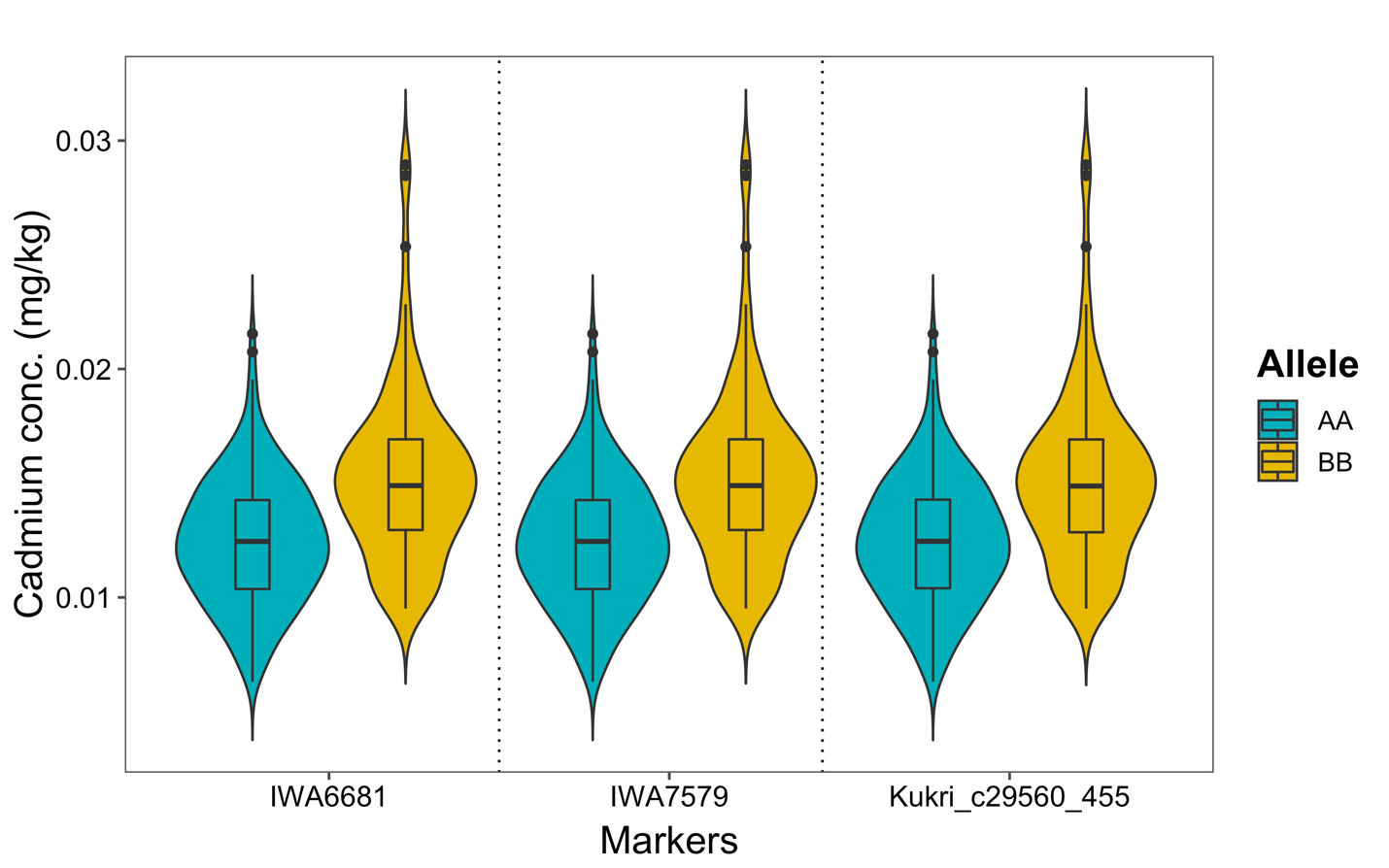

**Figure S5**: Violin plot showing the differences in the mean and variance of Cadmium concentration with alternative marker allele groups coded as AA and BB for the top three significant markers associated with mvQTL on chromosome 5A. Differences in both mean and variance can be observed across the marker groups indicating that these markers affect both mean and variance heterogeneity.
